# S-acylated Golga7b stabilises DHHC5 at the plasma membrane to regulate desmosome assembly and cell adhesion

**DOI:** 10.1101/481861

**Authors:** Keith T. Woodley, Mark O. Collins

## Abstract

S-acylation is the only fully reversible lipid modification of proteins however little is known about how protein S-acyltransferases (PATs) that mediate it are regulated. DHHC5 is a plasma membrane-localised PAT with roles in synaptic plasticity, massive endocytosis and cancer cell growth/invasion. Here we demonstrate that stabilisation of DHHC5 at the plasma membrane requires binding to and palmitoylation of an accessory protein Golga7b. This interaction requires the palmitoylation of the C-terminus of DHHC5 which regulates the internalisation of DHHC5 from the plasma membrane. Proteomic analysis of DHHC5/Golga7b-associated protein complexes reveals an enrichment in adhesion proteins, particularly components of desmosomes. We show that Desmoglein-2 and Plakophilin-3 are substrates of DHHC5 and that DHHC5/Golga7b are required for localisation of Desmoglein-2 to the plasma membrane and desmosomal patterning. Loss of DHHC5/Golga7b causes functional impairments in cell adhesion suggesting these proteins have a wider role in cell adhesion beyond desmosome assembly. This work uncovers a novel mechanism of DHHC5 regulation by Golga7b and demonstrates a role for the DHHC5/Golga7b complex in the regulation of cell adhesion.

## Introduction

S-acylation (palmitoylation) is unique amongst post-translational modifications as it is the only fully reversible modification that involves the attachment of a lipid to a protein, in this case, to the side chain of cysteine residues via a thioester bond. While palmitoylation was discovered several decades ago as a modification of viral proteins (Schmidt & Schlesinger, 1979), the progress in our understanding of palmitoylation has been slower than that for other modifications such as phosphorylation. For example, the regulation of the family of enzymes which attach the fatty acid onto proteins, palmitoyl acyltransferases (PATs) is still poorly understood. There are 23 PATs encoded in the human genome, all of which are integral membrane proteins (Korycka et al., 2012), with a range of cellular locations and tissue expression patterns (Ohno et al., 2006). They are also referred to as DHHC proteins as their catalytic tetrad contains the sequence Asp-His-His-Cys which is required for palmitoylation of substrates. However, it has been shown that in some cases, additional proteins are required for PAT activity. This was first demonstrated in yeast, where the palmitoylation of Ras, which is essential for its signalling and function (Hancock et al., 1990), requires a complex of two proteins localised to the endoplasmic reticulum, Erf2 and Erf4 (Lobo et al., 2002). Subsequently, it was discovered that in mammalian cells, DHHC9 requires the protein GCP16, also known as Golga7, to form a complex capable of palmitoylating Ras (Swarthout et al., 2005). The mechanism by which Erf4 acts on Erf2 was subsequently determined to involve stabilisation of the intermediate state where the acyl chain is transferred to the PAT before final transfer of the acyl chain to the substrate (Mitchell et al., 2012).

In a large scale affinity purification-mass spectrometry study, Golga7b was identified as a putative interactor of DHHC5 (Huttlin et al., 2015). Golga7b is a member of the Erf4 protein family which is closely related to GCP16 (Golga7) with 61% sequence identity. This interaction has yet to be confirmed and it is unknown whether Golga7b could regulate DHHC5 in a manner analogous to how DHHC9 is regulated by GCP16. Interestingly, Golga7b was also identified as a putative palmitoylated protein in both humans and mice (Morrison et al., 2015; Collins et al., 2017), although confirmation of this palmitoylation and the identity of the cognate PAT is lacking.

While the majority of PATs are localised to the Golgi, DHHC5 is localised to the plasma membrane (Ohno et al., 2006). It is expressed in all tissues in mammals but it is highly enriched in the heart and the brain. It plays a role in the growth and invasion of certain types of non-small-cell lung cancer and DHHC5 knockdown helps to prevent the growth and invasion of these types of tumours (Breusegem & Seaman, 2014). DHHC5 has few confirmed substrates, but it is known to palmitoylate the plasma membrane protein Flotillin-2, which is enriched in cholesterol rich microdomains (Li et al., 2012). In the heart, DHHC5 it is a key player in the phenomenon of massive palmitoylation-dependent endocytosis (MEND) (Hilgemann et al., 2013). This process occurs when oxygen is reintroduced to cardiac muscle cells starved of oxygen and involves the endocytosis of a large proportion of the plasma membrane, giving DHHC5 a role in the recovery of heart muscle from periods of anoxia (Lin et al., 2013). Additionally, an important role for DHHC5 in the regulation of the cardiac sodium pump through palmitoylation of phospholemman has been established (Howie et al., 2014).

In neurons, DHHC5 is present at the post-synaptic density and knockout mice have defects in learning and memory (Li et al., 2010). DHHC5 is retained at the plasma membrane of the post-synaptic density through interactions with the scaffolding protein PSD-95 and the tyrosine kinase Fyn (Brigidi et al., 2015). When neurons are activated by long-term potentiation, phosphorylation of DHHC5 by Fyn is reduced, DHHC5 is lost from the complex and is endocytosed to dendritic spines where it palmitoylates δ-catenin before both proteins are trafficked back to the plasma membrane at the post-synaptic density (Brigidi et al., 2015). This type of controlled endocytic movement of DHHC5 has not been described for any other PAT, but does leave a number of open questions including the mechanism of endocytosis from the plasma membrane and whether a similar system of trafficking of DHHC5 to and from the plasma membrane is present in other cell types or is unique to neurons.

We have identified a cluster of three palmitoylation sites in the C-terminus of DHHC5 (located just after its fourth transmembrane domain), some of which appear to be conserved in several other PATs (Collins et al., 2017). In brain extracts, DHHC exists in a predominantly palmitoylated form with several palmitoylated states apparent (Collins et al., 2017). The function of palmitoylation at these sites is unknown but we hypothesise that they may regulate DHHC5 localisation and/or stability. Indeed, recent work has established an important role for palmitoylation of the C-terminus of DHHC6 by DHHC16 in the regulation of its activity and turnover (Abrami et al., 2017). This region of DHHC5 is also involved in substrate recruitment; DHHC5 binds to and palmitoylates the sodium pump regulator phospholemman and this is dependent on a region of the C-terminus of DHHC5 just after its fourth transmembrane domain (Howie et al., 2014).

A number of studies have reported that several proteins involved in both cell:cell and cell:matrix adhesion are palmitoylated and that this palmitoylation regulates localisation and function of these proteins (Little et al., 1998; Lievens et al., 2016; Aramsangtienchai et al., 2017). Among adhesion protein complexes, desmosomes are notable as several components of the complex are palmitoylated (Roberts et al., 2014). Palmitoylation of plakophilin-3 is essential for both its inclusion into desmosomes and is required for the efficient assembly of desmosomes. One of the two desmosomal cadherins, desmoglein-2, is palmitoylated on a pair of cysteines (Roberts et al., 2016). Mutation of the palmitoylation sites of desmoglein-2 does not prevent incorporation of desmoglein-2 into desmosomes but does cause a trafficking defect with a portion of desmoglein-2 localised to lysosomes. This shows that it is likely that palmitoylation plays a central role in desmosomal assembly, turnover and trafficking however it is unknown which PAT or PATs regulate this process.

Here we demonstrate that Golga7b interacts with DHHC5 and Golga7b palmitoylation regulates plasma membrane localisation of DHHC5. We have determined that the interaction between DHHC5 and Golga7b is controlled by the palmitoylation of the C-terminal tail of DHHC5. We also show that palmitoylation of golga7b prevents internalisation of DHHC5 by AP2-regulated clathrin mediated endocytosis. Using affinity purification-mass spectrometry analysis we demonstrate that Golga7b defines the interactome of DHHC5 and contains known and novel substrates as well as regulators of DHHC5. Investigation of selected novel substrates revealed a central role for DHHC5 and Golga7b in cell adhesion; DHHC5 palmitoylates both desmoglein-2 and plakophilin-3 and loss of DHHC5 or Golga7b causes defects in cell adhesion. These results uncover a novel palmitoylation-dependent mechanism to control the localisation and function of DHHC5 and reveal an important new role for DHHC5 in cell adhesion.

## Results

### DHHC5 interacts with Golga7b in a palmitoylation-dependent manner

Initially, we set out to confirm whether Golga7b is indeed palmitoylated and if Golga7b interacts with DHHC5, as both of these results had come from large-scale studies and had not been validated by targeted methods (Huttlin et al., 2015; Collins et al., 2017). Initial attempts to express a palmitoylation-deficient mutant Golga7b in which cysteines in the sequence were mutated to alanines resulted in undetectable levels of expression by immunoblotting and immunofluorescence microscopy (data not shown). As palmitoylation regulates the stability of some proteins, this can lead to degradation of palmitoylation-deficient mutants (Ampah et al., 2018). Expression of mutant Golga7b with the addition of MG132 prevented degradation of mutant Golga7b (Fig. 1a) and therefore, we included MG132 in all subsequent experiments with this construct. To measure palmitoylation we used an acyl-biotin exchange (ABE) strategy (Drisdel et al., 2004) in which palmitoyl groups are selectively released by hydroxylamine at neutral pH, biotinylated and enriched using streptavidin resin. Using this approach, we found that a C-terminally FLAG-tagged full length wild-type (WT) Golga7b is palmitoylated when expressed in HeLa cells, but when the cysteines in the sequence were mutated to alanines, this palmitoylation was lost (Fig. 1a). Next, we sought to test whether DHHC5 could palmitoylate Golga7b using siRNA mediated knockdown of DHHC5 (Figure S1a) and detection of palmitoylation using ABE. When compared to cells that are treated with a non-targeting siRNA, there was a significant reduction in the palmitoylation state of Golga7b when DHHC5 is depleted (Fig. 1b) indicating that DHHC5 is the major PAT for Golga7b.

**Figure 1:**
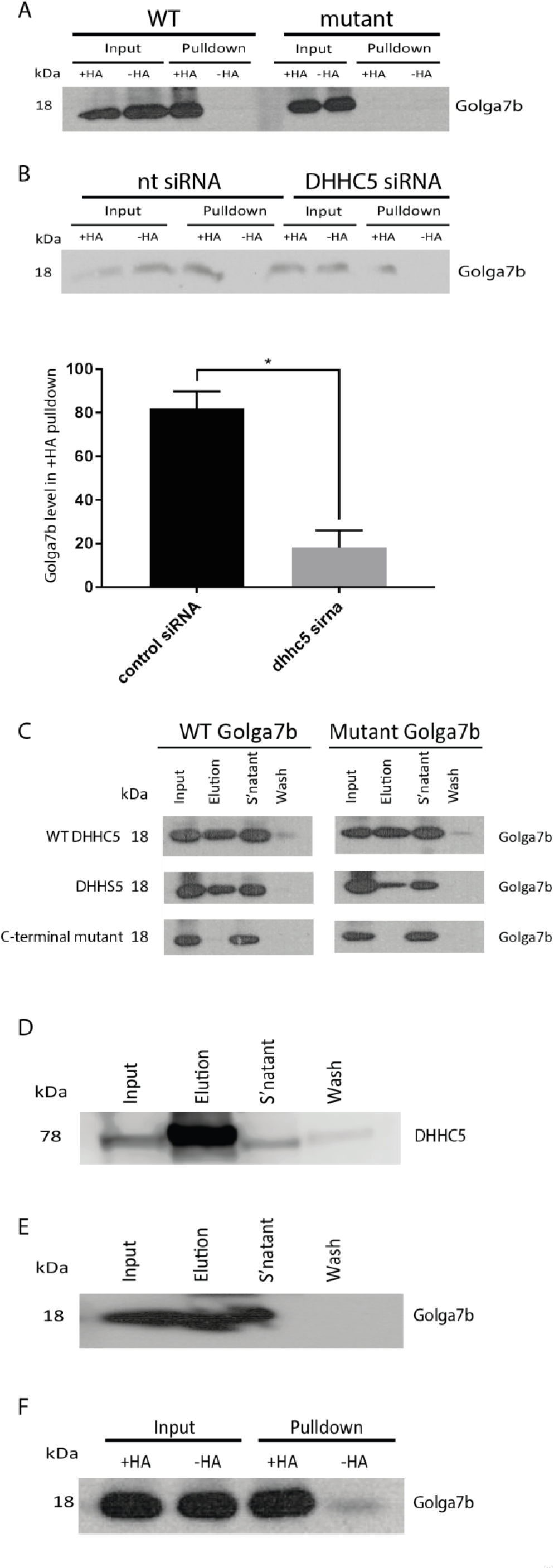
A palmitoylation-dependent interaction between DHHC5 and Golga7b. (A) ABE blot of WT Golga7b and a palmitoylation-deficient mutant form of the protein with the cysteines in the sequence replaced with alanine residues. The presence of Golga7b signal in the +HA lane of the WT pulldown and not in either the –HA lane or the +HA lane of the mutant shows that Golga7b is a palmitoylated protein. (B) ABE blot for Golga7b from HeLa cells treated with either DHHC5 siRNA or control non-targeting siRNA. A significant drop in the signal for Golga7b is seen in the +HA of the pulldown lane when DHHC5 is knocked down with siRNA. N=3, *=p<0.05, paired t-test. C – Co-IP blots of WT and non-palmitoylatable mutant Golga7b from pulldowns of WT DHHC5 and a pair of DHHC5 mutants (DHHS5 = catalytically inactive mutant where catalytic cysteine has been mutated to serine, C-terminal mutant = mutant where the 3 C-terminal palmitoylation sites of DHHC5 have been mutated to alanine residues.). WT and DHHS5 could pulldown both WT and mutant Golga7b but the C-terminal mutant was unable to pull down either form of Golga7b showing that the palmitoylation of this region of DHHC5 is essential for the stability of this interaction. (D) Co-IP of DHHC5 with Golga7b from whole mouse forebrain extract. (E) Co-IP of Golga7b with DHHC5 from whole mouse forebrain extract. (F) ABE assay of Golga7b from mouse forebrain extract showing a hydroxylamine sensitive signal for Golga7b indication confirming that it is palmitoylated in mouse forebrain extract.

We attempted to confirm the interaction between DHHC5 and Golga7b by co-immunoprecipitation using N-terminally HA-tagged DHHC5 and FLAG-tagged WT and palmitoylation-deficient mutant Golga7b constructs co-expressed in HeLa cells. DHHC5 immunoprecipitated both WT and mutant Golga7b (Fig. 1c) which indicates that the palmitoylation state of Golga7b does not affect its interaction with DHHC5. The same is also true for a catalytically inactive form of DHHC5 where the catalytic cysteine is mutated to a serine residue (DHHS5, Figure 1c). However, when the three palmitoylation sites present in the C-terminal tail of DHHC5 (Collins et al., 2017) are mutated, this mutant DHHC5 was unable to immunoprecipitate either the WT or mutant Golga7b (Figure 1c) suggesting that palmitoylation of DHHC5 is directly required for this interaction. To confirm that the interaction of DHHC5 and Golga7b occurs *in vivo*, we performed immunoprecipitations of endogenous DHHC5 and Golga7b from a mouse forebrain lysate. This demonstrated that DHHC5 can be immunoprecipitated by endogenous Golga7b (Figure 1d) and that endogenous DHHC5 can reciprocally immunoprecipitate Golga7b (Figure 1e). We next confirmed that endogenous Golga7b is palmitoylated in this context using an ABE assay (Figure 1f). These results demonstrate that DHHC5 and Golga7b interact *in vivo* and that DHHC5 is the major PAT for Golga7b.

### Co-expression of Golga7b and DHHC5 drives the complex to the plasma membrane

We confirmed that DHHC5 is localised to the plasma membrane (Ohno et al., 2006) by staining for the endogenous protein in HeLa cells (Figure S2a). It has been reported that the over-expression of PATs can lead to their mislocalisation (Hou et al., 2009), which could suggest that factors that control their location need to be present in sufficient quantities to ensure proper localisation. When DHHC5 is expressed alone, staining is seen throughout the cell (Figure S2b) indicating that indeed mislocalisation occurs upon overexpression. We hypothesised that Golga7b might act as a regulator of DHHC5 and may be involved in the targeting or stabilisation of DHHC5 at the plasma membrane. To investigate this, we co-expressed DHHC5 with either WT or a palmitoylation-deficient mutant Golga7b and performed immunofluorescence microscopy. When DHHC5 is co-expressed with WT Golga7b, the majority of DHHC5 is present at the plasma membrane (Figure 2a). This indicates that when DHHC5 is overexpressed alone, there is insufficient endogenous Golga7b present to target or stabilise all DHHC5 protein to the plasma membrane. Strikingly, when DHHC5 is co-expressed with mutant Golga7b which cannot be palmitoylated, the majority of DHHC5 is lost from the plasma membrane (Figure 2b). This indicates that palmitoylation of Golga7b is required for the maintenance of DHHC5 at the plasma membrane and when this is prevented, DHHC5 is mislocalised.

**Figure 2:**
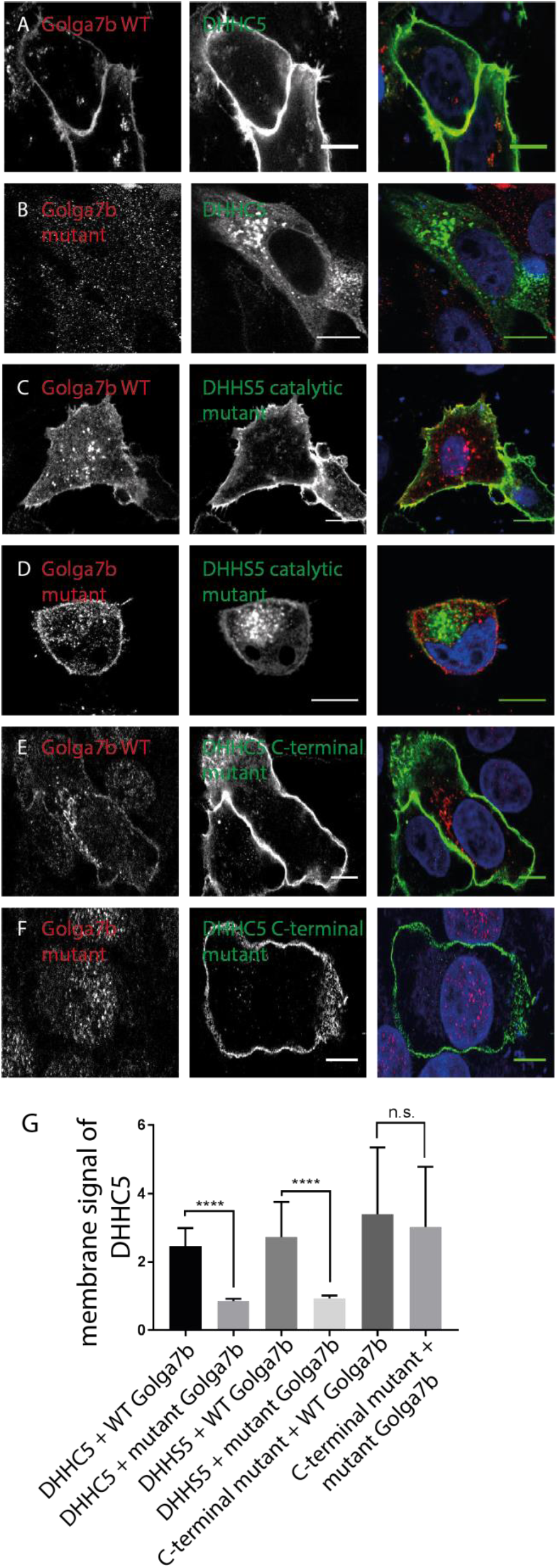
Regulation of DHHC5 localisation by palmitoylation of Golga7b and the C-terminus of DHHC5. (A) Confocal images of HeLa cells co-expressing WT Golga7b (1st column) and DHHC5 (2nd column), with a merged image (3rd column). (B) – confocal images of HeLa cells co-expressing mutant Golga7b (1st column) and DHHC5 (2nd column), with a merged image (3rd column). (C) Confocal images of HeLa cells co-expressing WT Golga7b (1st column) and DHHS5 (2nd column), with a merged image (3rd column). (D) Confocal images of HeLa cells co-expressing mutant Golga7b (1st column) and DHHS5 (2nd column), with a merged image (3rd column). (E) Confocal images of HeLa cells co-expressing WT Golga7b (1st column) and DHHC5 C-terminal mutant (2nd column), with a merged image (3rd column). (F) Confocal images of HeLa cells co-expressing mutant Golga7b (1st column) and DHHC5 C-terminal mutant (2nd column), with a merged image (3rd column). (G) Quantification of the membrane signal of DHHC5 in each of the images (A-F). ****=p<0.001, n.s. = not significant, unpaired t-test. N=49, DHHC5 + WT Golga7b. N=53, DHHC5 + mutant Golga7b. N=40, DHHS5 + WT Golga7b. N=28, DHHS5 + Golga7b mutant. N=42, C-terminal mutant + WT Golga7b. N=31, C-terminal mutant + mutant Golga7b.

Co-expression of wild-type and mutant Golga7b with the catalytic mutant of DHHC5 had the same effect on the localisation of DHHS5 as it does on the WT DHHC5 (Figures 2c and d), and over-expression of DHHS5 alone again shows a similar mislocalisation as seen with wild-type DHHC5 (Figure S2c). This would initially suggest that DHHC5 does not need to be enzymatically active for Golga7b to target DHHC5 to the plasma membrane. However, given that there is sufficient endogenous DHHC5 activity to palmitoylate overexpressed Golga7b (Figure 1a), this is unlikely to be the case. Next, we sought to investigate the role of palmitoylation at the C-terminus of DHHC5 and its effect on DHHC5 localisation given that the palmitoylation-deficient mutant DHHC5 is unable to interact with Golga7b. Unexpectedly, when we co-expressed the DHHC5 C-terminal palmitoylation-deficient mutant with either wild-type or mutant Golga7b (Figure 2e and f) it was localised to the plasma membrane. Furthermore, it was localised to the plasma membrane when over-expressed alone without Golga7b (Figure S2d).

### Palmitoylation of Golga7b regulates localisation of endogenous DHHC5 to the plasma membrane

In order to demonstrate that Golga7b regulates the localisation of endogenous DHHC5, we expressed WT or mutant Golga7b in HeLa cells and used immunofluorescence microscopy to probe the localisation of endogenous DHHC5 (Figure 3). Expression of WT Golga7b significantly increased levels of endogenous DHHC5 at the plasma membrane compared to endogenous DHHC5 without Golga7b over-expression (Figure 3a, 3c and f) indicating that the expression level of Golga7b regulates the amount of plasma membrane localised DHHC5. Interestingly, over-expression of mutant Golga7b reduces endogenous DHHC5 localisation at the plasma membrane (Figure 3b and f), similar to what we observed when both DHHC5 and mutant Golga7b are over-expressed. This indicates that expression of a palmitoylation-deficient form of Golga7b prevents stabilisation of DHHC5 at the plasma membrane in a dominant negative manner.

**Figure 3:**
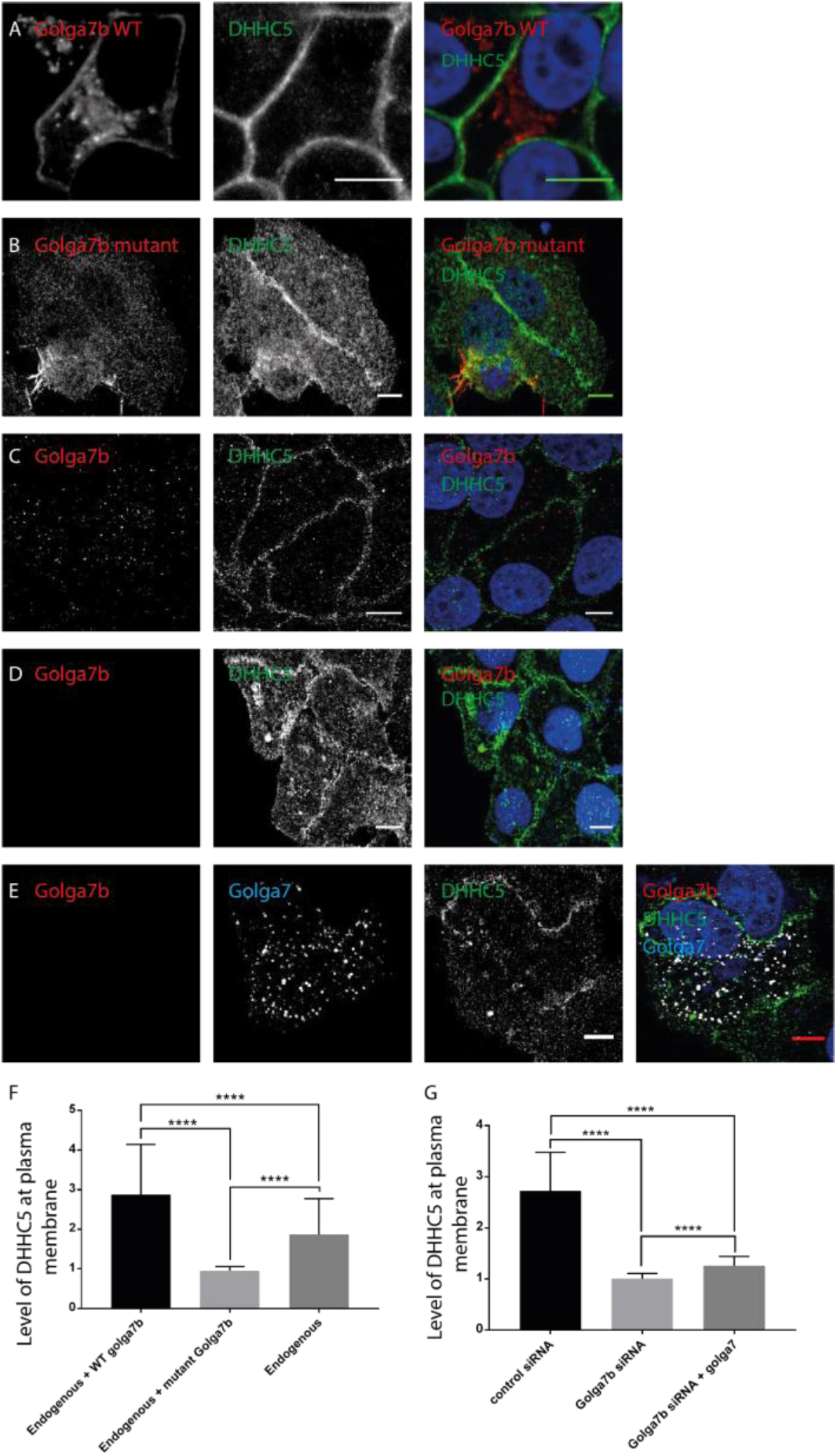
Regulation of plasma membrane localisation of endogenous DHHC5 by expression of Golga7b. (A) Confocal images of endogenous DHHC5 in HeLa cells over-expressing mutant Golga7b. (B) Confocal images of endogenous DHHC5 in HeLa cells over-expressing WT Golga7b. (C) Confocal images of endogenous Golga7b and DHHC5 in HeLa cells treated with negative control non-targeting siRNA. (D) Confocal images of endogenous Golga7b and DHHC5 in HeLa cell treated with Golga7b siRNA. (E) Confocal images of endogenous Golga7b and DHHC5 in HeLa cell treated with Golga7b siRNA and expressing Golga7. (F) Quantification of plasma membrane signal of endogenous DHHC5 when co-expressed with WT or mutant Golga7b or without any transfected proteins. ****=P<0.001, t-test, endogenous + WT Golga7b n=37, endogenous + mutant Golga7b n=23, endogenous n=42. (G) Quantification of the plasma membrane signal of DHHC5 in A-C, ****=p<0.001 t-test, control siRNA n=23, Golga7b siRNA n=23, Golga7b siRNA + Golga7 n=27.

In order to more definitively show that insufficient Golga7b leads to DHHC5 mislocalisation as was suggested by over-expression of DHHC5 alone (Figure S2b), we depleted Golga7b with siRNA in HeLa cells (Figure 3d). We observed that the localisation of endogenous DHHC5 was significantly reduced at the plasma membrane upon Golga7b depletion, compared to cells treated with a non-targeting negative control siRNA where DHHC5 was present almost exclusively at the plasma membrane (Figure 3c and d). This confirms that sufficient levels of Golga7b are required to localise DHHC5 to the plasma membrane. To ensure this was an effect specific to Golga7b, we attempted to rescue the localisation defect of DHHC5 due to Golga7b depletion by expression of the closely related Golga7 (GCP16) (Figure 3e). Over-expression of Golga7 in Golga7b depleted cells failed to completely restore DHHC5 to the plasma membrane, suggesting that this effect is specific to Golga7b and cannot be significantly rescued by related proteins (Figure 3g).

### Golga7b defines DHHC5 interactomes

As Golga7b has such a profound effect on the localisation of DHHC5, we used this to investigate whether the interactions of DHHC5 are different when it is present at the plasma membrane compared to when it is localised in endomembranes. To achieve this, a C-terminally triple FLAG-tagged DHHC5 was co-expressed with either WT or mutant Golga7b in HeLa cells before performing immunoprecipitations. Initially, we sought to prove that DHHC5 would immunoprecipitate proteins differentially when present at the plasma membrane or within the cell. We chose Flotillin-2 to test this as it is a known DHHC5 substrate (Li et al., 2012) and is present at the plasma membrane. We showed that DHHC5 was able to immunoprecipitate significantly more Flotillin-2 when it was present at the plasma membrane (co-expressed with WT Golga7b) compared to when it was present at endomembranes (co-expressed with palmitoylation-deficient mutant Golga7b) (Figure 4a).

**Figure 4:**
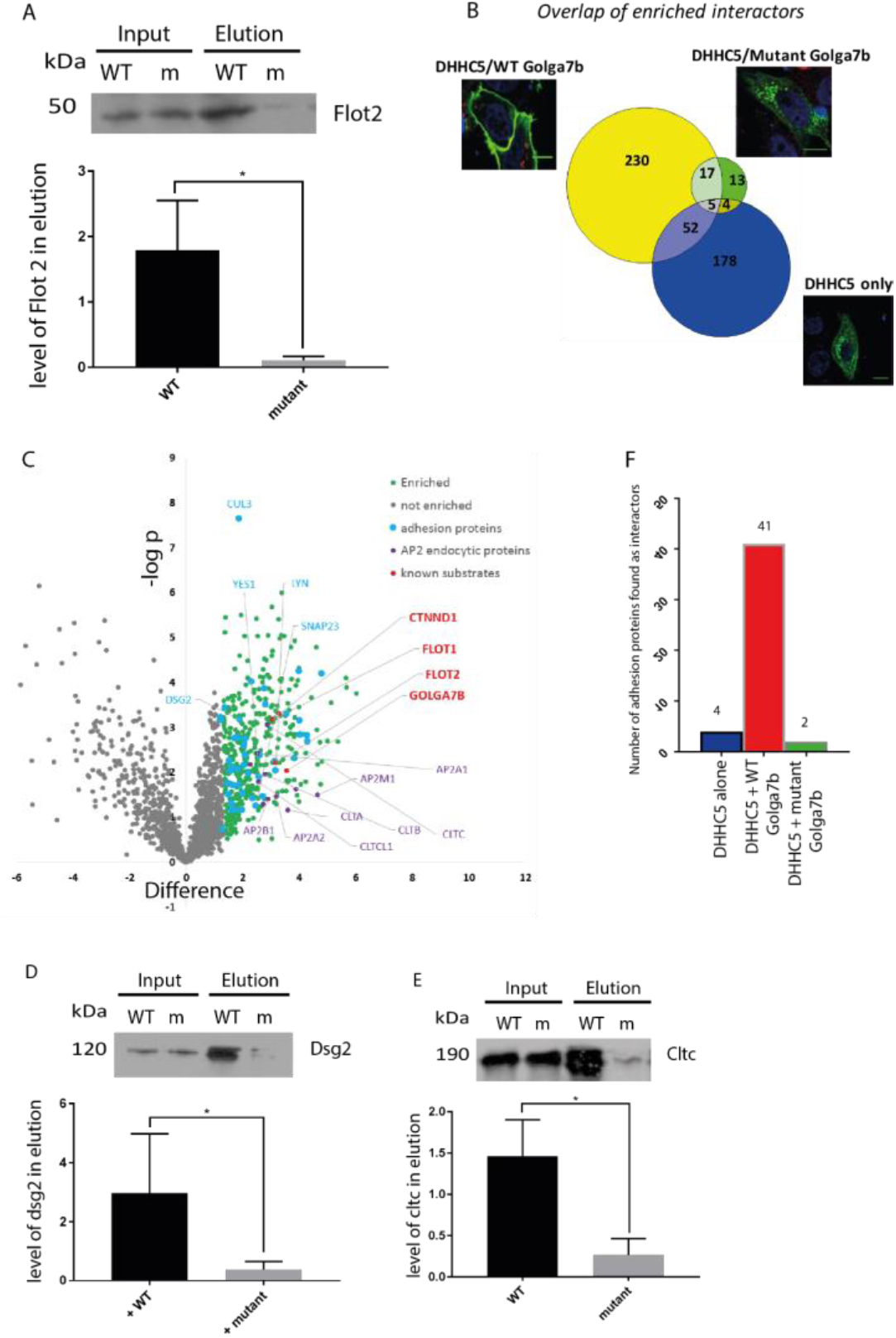
Proteomic analysis of context-dependent DHHC5-associated protein complexes. (A) Co-IP western blots of Flotillin-2 from Hela cells co-expressing DHHC5 and either wild-type (WT) or mutant (m) Golga7b and quantification normalised to input, *=p<0.05, ratio paired t-test, n=3. (B) Venn diagram comparing the overlap of numbers of proteins immunoprecipitated when DHHC5 is expressed alone or co-expressed with either WT or mutant Golga7b. (C) Scatter plot of DHHC5 interactors from AP-MS analysis from HeLa cells co-expressing DHHC5 and WT Golga7b highlighting interacting proteins involved in clathrin-mediated endocytosis, cell:cell adhesion and known substrates of DHHC5. (D) Co-IP western blots of desmoglein-2 from Hela cells co-expressing DHHC5 and either Wwild-type (WT) or mutant (m) Golga7b and quantification normalised to input, *=p<0.05, ratio paired t-test, n=3. (E) Co-IP western blots of clathrin heavy chain from Hela cells co-expressing DHHC5 and either wild-type (WT) or mutant (m) Golga7b and quantification normalised to input, *=p<0.05, ratio paired t-test, n=3. (F) Bar graph comparing the number of the cell:cell adhesion proteins identified in each protein interaction dataset.

In order to explore this more widely, we used an affinity purification-mass spectrometry (AP-MS) approach in which DHHC5 complexes were purified and competitively eluted by triple-FLAG peptide, separated by SDS PAGE, subjected to in-gel digestion and analysed by quantitative tandem mass spectrometry. We defined sets of DHHC5 interactors based on their quantitative enrichment in IPs versus control (non-transfected) purifications when a) DHHC5 was localised to the plasma membrane (co-expressed with WT Golga7b), b) DHHC5 was localised at endomembranes (co-expressed with palmitoylation-deficient mutant Golga7b), c) DHHC5 was localised at endomembranes (expressed alone). In order to rule out any effects of addition of MG132 when we co expressed DHHC5 with palmitoylation-deficient mutant Golga7b, we also generated and analysed IPs from cells which we co-expressed DHHC5 with wild-type Golga7b +/-MG132. Comparison of enriched proteins in these IPs +/-MG132 identified only one protein differentially enriched by the addition of MG132 demonstrating that it did not significantly affect the set of DHHC5 interactors.

Data from these experiments revealed that DHHC5 interacts with many more proteins when localised at the plasma membrane by wild-type Golga7b (304 proteins, Figure 4b, Table S1) compared to when it is expressed with mutant Golga7b when it is localisaed to endomembranes (39 proteins, Table S2), with a small number of proteins common to both conditions (22 proteins) (t-test with a permutation based FDR 0.05). DHHC5 does interact with a large number of proteins at the plasma membrane (304 proteins when DHHC5 is co-expressed with WT Golga7b), however, this is not uncommon for enzymes which may have a large number of substrates. When DHHC5 is expressed alone (Table S3) it interacts with some proteins also identified when it was co-expressed with WT Golga7b (52 proteins). Among these are members of the AP2 complex with is involved in certain types of clathrin-mediated endocytosis and has been shown to be involved in endocytosis of DHHC5 (Brigidi et al., 2015). Among the proteins that are specific to the dataset obtained when DHHC5 is expressed on its own, perhaps the most interesting is a large number of proteins involved with the proteasome and protein degradation pathways (35 proteins). These proteins are not present when DHHC5 is expressed with either WT or mutant Golga7b, suggesting that Golga7b may offer DHHC5 protection from internalisation and degradation by stabilising DHHC5 at the plasma membrane. When DHHC5 is expressed alone there are many more cytoplasmic proteins identified as interactors compared to when it is co-expressed with WT Golga7b; this is likely caused by the mislocalisation of DHHC5 in the absence of exogenous Golga7b.

When we directly compared the interaction datasets from DHHC5 co-expressed with WT or mutant Golga7b, a number of proteins were differentially enriched; nine were enriched in the DHHC5/mutant Golga7b dataset, and 163 in the DHHC5/WT Golga7b dataset (t-test with a permutation based FDR 0.05). This clearly demonstrates the profound effect of Golga7b-dependent localisation of DHHC5 has on its interactome. To validate this, we chose two proteins that were differently enriched between the WT and mutant Golga7b conditions that were not known DHHC5 interactors or substrates. Both desmoglein-2, a desmosomal cadherin, and the clathrin heavy chain, an important endocytosis protein, interacted with DHHC5 significantly more when DHHC5 was at the plasma membrane (co-expressed with WT Golga7b) compared to when it was at endomembranes (co-expressed with mutant Golga7b) (Figures 5d and e), showing that both of these proteins are bona fide DHHC5 interactors and that our AP-MS results are reliable.

**Figure 5:**
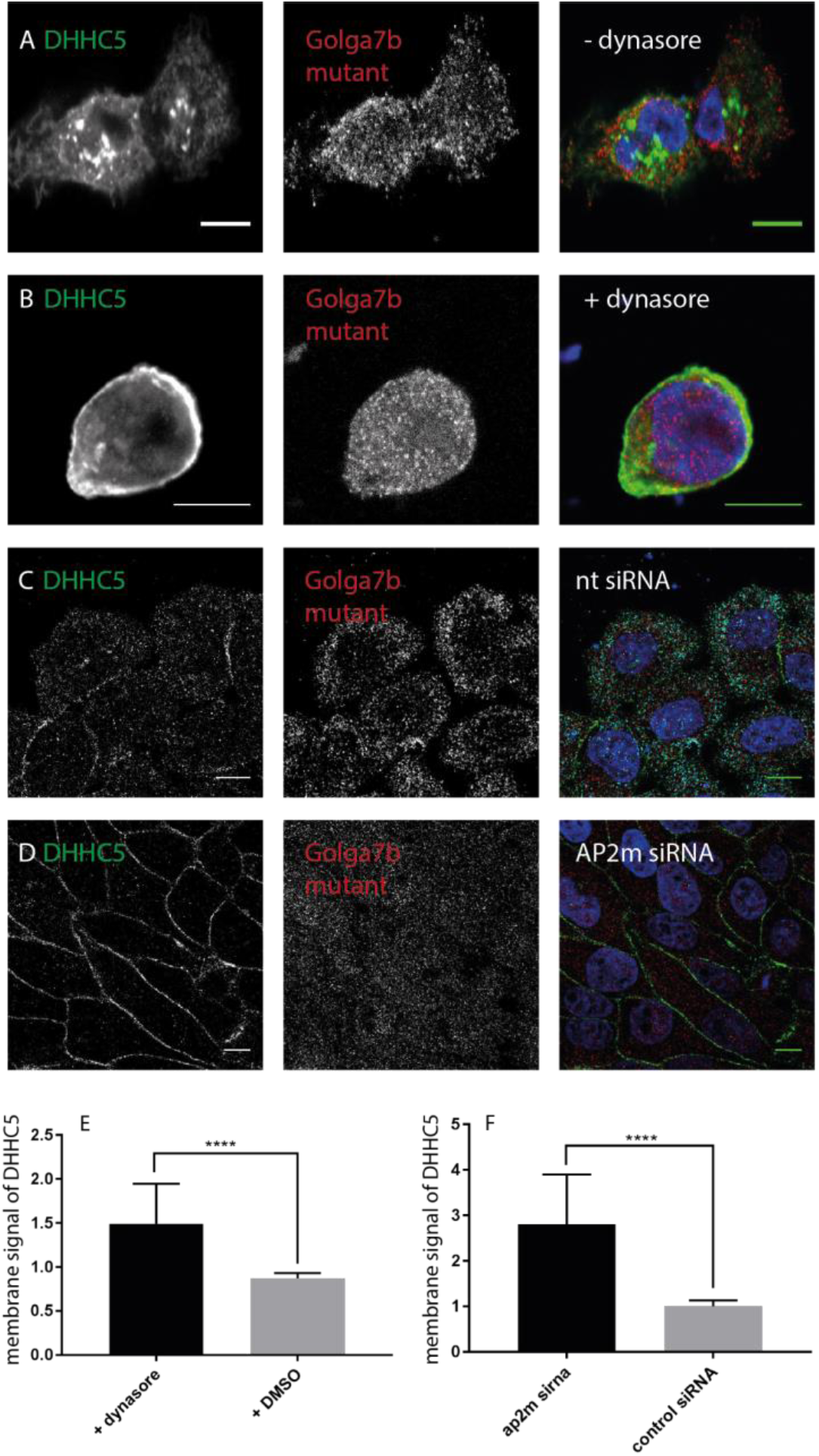
AP2-mediated endocytosis of DHHC5 requires palmitoylated Golga7b. (A) 30 minute DMSO treatment of HeLa cells co-expressing WT DHHC5 and mutant Golga7b. (B) 30 minute Dynasore treatment (25µM) of HeLa cells co-expressing WT DHHC5 and mutant Golga7b. (C) Non-targeting control siRNA treatment (10nM) of HeLa cells expressing mutant Golga7b. (D) AP2µ siRNA treatment (10nM) of HeLa cells that were subsequently transfected with mutant Golga7b. (E) Quantification of plasma membrane signal of DHHC5 in A and B. ****=p<0.001, unpaired t-test. N=37, +dynasore. N=28, +DMSO. (F) Quantification of plasma membrane signal of DHHC5 in D and E. ****=p<0.001, unpaired t-test.

Among the set of proteins that were significantly enriched when DHHC5 was expressed with WT Golga7b compared to mutant Golga7b was a number of proteins associated with clathrin-mediated endocytosis, including three major members of the AP2 complex (Figure 4c), plus the clathrin light and heavy chain. This indicates that the endocytosis of DHHC5 may be mediated by this complex, as DHHC5 is known to be actively trafficked away from the plasma membrane in neurons (Brigidi et al., 2015) and has a number of endocytic sequences in its C-terminal tail. When looking at the proteins that interact with DHHC5 when it is co-expressed with WT Golga7b, nine of the top ten over-represented gene ontology terms are involved with cell:cell or cell:matrix adhesion (Figure 4f). While it is known that a number of proteins involved in adhesion are palmitoylated (Roberts et al., 2014; Lievens et al., 2016; Aramsangtienchai et al., 2017), the PATs responsible are mostly unknown. Given the large number of proteins immunoprecipitated by DHHC5, particularly in the desmosomal cell adhesion complex, we hypothesised that DHHC5 is the PAT responsible for the palmitoylation of these proteins and could be a key player in regulating cellular adhesion. In particular, it was noted that the desmosomal cadherin Desmoglein-2, which is a known palmitoylated protein (Roberts et al., 2014), and the Tight junction protein ZO-1 which opens up the possibility that palmitoylation by DHHC5 is an important regulatory process in these two different processes in cell adhesion.

### Palmitoylation of golga7b prevents internalisation of DHHC5

As several of the clathrin mediated endocytosis complex members were found as interactors of DHHC5 in our AP-MS study (Figure 4c), we decided to further investigate the mechanism of DHHC5 internalisation and endocytosis. Given that palmitoylation of Golga7b is required to localise DHHC5 at the plasma membrane, we sought to better understand the mechanism of internalisation of DHHC5. We first investigated whether it was possible to rescue mislocalisation of DHHC5 when co-expressed with mutant Golga7b by inhibiting the clathrin-mediated endocytosis. When HeLa cells expressing WT DHHC5 and mutant Golga7b are treated with 25µM dynasore, the majority of DHHC5 is stabilised at the plasma membrane after a relatively short time (within 30 minutes) (Figure 5a and b). This shows that DHHC5 is rapidly trafficked to and from the plasma membrane in a number of cell types outside of what has been found previously in neurons (Brigidi et al., 2015). It also suggests a role for Golga7b in stabilising DHHC5 at the plasma membrane by preventing endocytosis.

As dynasore may have some off-target effects (Park et al., 2013), we validated these observations using siRNA knockdown of the AP2µ subunit, which was found as an interactor of DHHC5 in the AP-MS experiments. When AP2µ is depleted, it leads to a significant drop in the endocytosis of the transferrin receptor and had a noticeable impact on the number of clathrin coated pits formed in cells (Motley et al., 2003). When we depleted AP2µ in HeLa cells co-expressing WT DHHC5 and mutant Golga7b, DHHC5 was restored to the plasma membrane, while in cells treated with a control non-targeting siRNA DHHC5 remained within the cell (Figure 5c and d). This confirms that DHHC5 requires AP2µ for endocytosis from the plasma membrane (Brigidi et al., 2015) and that palmitoylation of Golga7b prevents endocytosis of DHHC5. Importantly, this demonstrates conclusively that palmitoylation deficient Golga7b does not prevent DHHC5 association with the plasma membrane and that palmitoylated Golga7b acts by stabilising DHHC5 at the plasma membrane.

### DHHC5/Golga7b regulates desmosome trafficking & assembly

As a number of proteins involved in cell adhesion were found as DHHC5 interactors in our AP-MS study (Figure 4c), we wanted to investigate whether any of these proteins were DHHC5 substrates and if DHHC5 is involved in cell adhesion. Among these is the important desmosomal cadherin desmoglein-2 (DSG2, Figure 4d). DSG2 is a known palmitoylated protein (Roberts et al., 2016), where its palmitoylation regulates the localisation and trafficking of the protein to the plasma membrane. Because of this, we decided to investigate whether DHHC5 is the PAT responsible for DSG2 palmitoylation, so we performed an ABE assay on DHHC5 depleted HeLa cells and control siRNA treated cells (Figure 6a). We found that upon DHHC5 depletion, DSG2 palmitoylation is almost completely abolished, showing that DHHC5 is the PAT responsible for the majority, if not all, of DSG2 palmitoylation. It also suggests that our AP-MS approach allows the discovery of new substrates of PATs, however, this may not be viable in all cases as it is known that some PATs only interact weakly with some of their substrates (Lemonidis et al., 2014).

**Figure 6:**
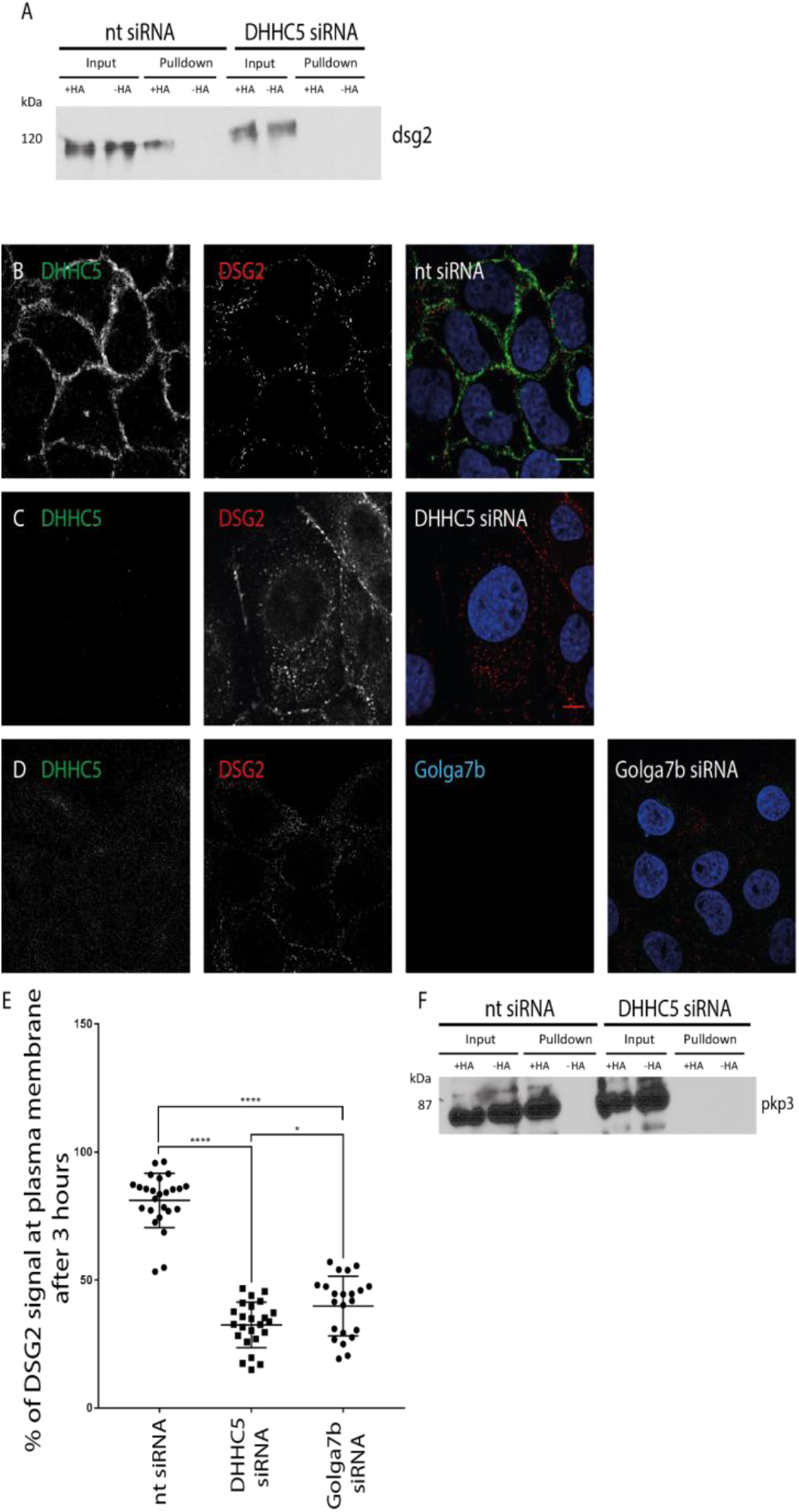
Desmogelin-2 palmitoylation by DHHC5/Golga7b regulates its plasma membrane localisation and incorporation into desmosomes. (A) ABE western blot for desmoglein-2 (DSG2) from HeLa cells treated with 10nM non-targeting siRNA (nt siRNA) or DHHC5 siRNA. Loss of the band in the +HA pulldown in the DHHC5 siRNA condition shows that DSG2 palmitoylation is dependent on DHHC5. (B) Confocal images of DHHC5 and DSG2 in nt siRNA treated A431 cells three hours after calcium was reintroduced to the media in calcium switch experiments. (C) Confocal images of DHHC5 and DSG2 in DHHC5 siRNA treated A431 cells three hours after calcium was reintroduced to the media in calcium switch experiments. (D) Confocal images of DHHC5, DSG2 and Golga7b in Golga7b siRNA treated A431 cells three hours after calcium was reintroduced to the media in calcium switch experiments. (E) Quantification of the percentage of total signal DSG2 seen at the plasma membrane and within the cell (internal) after 3 hours in calcium switch experiments in either nt siRNA or DHHC5 siRNA conditions. ****=p<0.001, unpaired t-test. N=24, DHHC5 siRNA. N=25, control siRNA. N=26, Golga7b siRNA. (F) ABE western blot for plakophilin-3 (pkp3) from HeLa cells treated with 10nM non-targeting siRNA (nt siRNA) or DHHC5 siRNA. Loss of the band in the +HA pulldown in the DHHC5 siRNA condition shows a DHHC5-dependent loss of pkp3 palmitoylation.

Given that palmitoylation of DSG2 regulates its localisation to the plasma membrane and trafficking it to desmosomes, we investigated the effect of DHHC5 knockdown on the localisation and trafficking of DSG2 using immunofluorescence microscopy of A431 cells with DHHC5 depletion and calcium switch assays. In culture, desmosomes are calcium dependent, so when cells are cultured in a medium that is free from calcium desmosomes are rapidly internalised and trafficked back to the plasma membrane when calcium is re-introduced. In A431 cells treated with a non-targeting negative control siRNA, DSG2 returned to the plasma membrane gradually over the three-hour time course when calcium was added to the media (Figure 6b, Figure S3). However, when DHHC5 was depleted, DSG2 signal remained within the cell, with no significant change in the level of DSG2 at the plasma membrane during the three-hour time course (Figure 6c, Figure S4). This shows that depletion of DHHC5 significantly inhibits the trafficking of DSG2 to the plasma membrane.

In order to establish if Golga7b plays a functional role in desmosome assembly through its regulation of DHHC5 localisation, we determined the effect of Golga7b depletion on the trafficking of DSG2. As with DHHC5, A431 cells were treated with Golga7b siRNA and a calcium switch performed. Again, after three hours of calcium being returned to the media, DSG2 had not returned to the plasma membrane at the same level as cells treated with a non-targeting siRNA (Figure 6d). This links DHHC5 location to the correct trafficking of DSG2 and implicates Golga7b in the regulation of DHHC5 substrates.

The DSG2 localisation phenotype is similar to that seen when palmitoylation of the desmosomal protein Plakophilin-3 (PKP3) is prevented (Roberts et al., 2014). PKP3 was not found as an interactor of DHHC5, but the localisation of DSG2 after DHHC5 depletion is similar to palmitoylation-deficient PKP3 mutants (Roberts et al., 2014). We treated HeLa cells with either DHHC5 siRNA or non-targeting control siRNA, performed ABE assays to measure palmitoylation levels of PKP3 and found that there was a significant and almost total loss of palmitoylation of PKP3 compared to a non-targeting siRNA (Figure 6f). This shows that PKP3 is a DHHC5 substrate and makes it likely that the phenotype seen with DSG2 localisation and trafficking is caused by a combination of the loss of palmitoylation of both DSG2 and PKP3.

### DHHC5/Golga7b regulates cell adhesion

To investigate whether DHHC5 plays a more general role in cell adhesion, or if the localisation and palmitoylation defects observed in desmosomes are sufficient to cause a loss of cell adhesion, we used a modified form of a cell scatter assay. When cells are exposed to a growth factor they internalise their cell:cell and cell:matrix adhesion proteins which cause cells to scatter away from the colonies they naturally form and mimics epithelial to mesenchymal transition. All cell adhesion proteins are targeted in this assay, but a deficiency in one of more aspects of the adhesion machinery will lead to a change in scattering. The speed and distance of this scattering are dependent on cell adhesion with reduced adhesion leading to enhanced scattering. When A431 cells were subject to DHHC5 depletion before treatment with low levels of EGF (5ng/ml), cells scattered noticeably (Figure 7b and c), whereas control non-targeting siRNA treated cells remained in colonies when treated with the same concentration of EGF (Figure 7a and c). This shows that depletion of DHHC5 leads to a reduction of adhesion of cells to one another and implicates DHHC5 in a wider role in cell:cell adhesion, as in order to scatter, cells must lose all cell:cell junctions including desmosomes and tight junctions (Chen, 2005).

**Figure 7:**
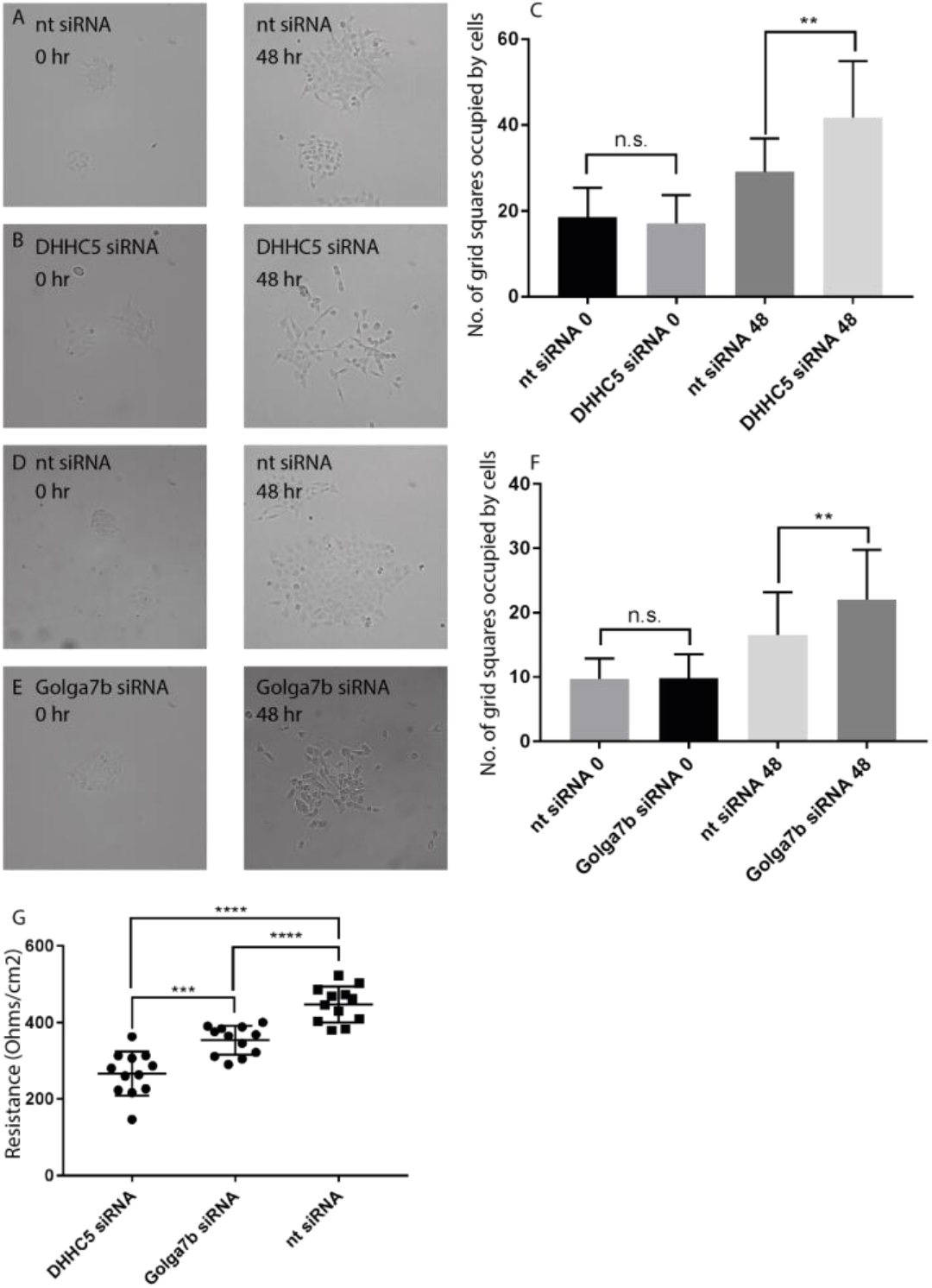
Cell adhesion is regulated by DHHC5/Golga7b. (A) 10x brightfield images of A431 cells treated with non-targeting siRNA at time 0 (left) and after 48 hours (right) of 5 ng/ml EGF treatment. (B) 10x brightfield images of A431 cells treated with DHHC5 siRNA at time 0 (left) and after 48 hours (right) of 5ng/ml EGF treatment. (C) Quantification of cell scattering of A431 cells, showing a significant difference in cell scatter after 48 hours of DHHC5 knockdown. N=3, n=31. **= p<0.005, unpaired t-test. (D) 10x brightfield images of A431 cells treated with non-targeting siRNA at time 0 (left) and after 48 hours (right) of 5ng/ml EGF treatment. (E) 10x brightfield images of A431 cells treated with Golga7b siRNA at time 0 (left) and after 48 hours (right) of 5ng/ml EGF treatment. (F) Quantification of cell scattering of A431 cells, showing a significant difference in cell scatter after 48 hours of DHHC5 knockdown. N=3, n=31. **= p<0.005, unpaired t-test. (G) Plot showing the resistance of A431 cells from a trans-epithelial electrical resistance assay treated with either DHHC5 siRNA, Golga7b siRNA or non-targeting siRNA. N=12, ****=p<0.001, unpaired t-test, ***=p<0.01, unpaired t-test

To test if depletion of Golga7b mirrors the effects seen with DHHC5 depletion in the calcium switch assay, we performed cell scatter assays on Golga7b depleted A431 cells (Figure 7 d-f). We observed enhanced scattering of cells upon Golga7b depletion compared to control cells exposed to a non-targeting siRNA. This shows Golga7b has a functional role in cell adhesion which is likely to be caused by its regulation of DHHC5 localisation. Given that we have shown that DHHC5 interacts not only with DSG2 but also with Tight junction protein ZO-1 (Table S2), it is possible that the wider effects that DHHC5 depletion has on cell adhesion could be caused by disruption of both desmosomes and tight junctions.

To further demonstrate a functional role of DHHC5 and Golga7b in cell adhesion, we performed trans-epithelial electrical resistance assays. This involves the growth of cells on a semi-permeable membrane support and measurement of the electrical resistance across this membrane (Chen et al., 2015); the stronger the cell adhesion, the higher the resistance measurement will be. When we performed this assay on confluent A431 cells that were treated with either DHHC5 or negative control non-targeting siRNA, we observed a significant reduction in electrical resistance in cell cultures treated with DHHC5 or Golga7b siRNA (Figure 7g). This shows that the adhesion between the cells is reduced in these conditions highlighting that the DHHC5/Golga7b complex is an important regulator of the cell adhesion machinery.

## Discussion

The regulation of subcellular localisation of many PATs is poorly understood, and DHHC5 is unique among them as it is the only known PAT to exhibit dynamic changes in localisation in response to external stimuli (Brigidi et al., 2015). While it has been shown in neurons that the localisation of DHHC5 is regulated by phosphorylation, we have shown that the localisation of DHHC5 is also regulated by the interaction and palmitoylation of an accessory protein, Golga7b. It is notable that the over-expression of palmitoylation-deficient mutant Golga7b causes mislocalisation of endogenous DHHC5 which suggests that Golga7b has a dominant negative effect on DHHC5. Over-expression of the WT Golga7b increased localisation of endogenous DHHC5 at the plasma membrane, which further strengthens the argument that the level of Golga7b within the cell is an important determinant of DHHC5 localisation. The catalytically dead DHHC5 (DHHS5) mutant like wild-type DHHC5 localises to the plasma membrane when co-expressed with Golga7b. However, wild-type Golga7b is readily palmitoylated when expressed in HeLa cells (Figure 1a), which suggests that endogenous DHHC5 is sufficient to palmitoylate Golga7b to a level required for correct localisation of DHHS5.

Palmitoylation sites in the C-terminus of DHHC5 are required for its interaction with Golga7b and stabilisation at the plasma membrane. Mutation of these palmitoylation sites appears to prevent internalisation of DHHC5 from the plasma membrane. Sorting signals in the C-terminal tails of PATs are important factors in controlling their localisation and are able to switch the localisation of a PAT from the Golgi to the ER (Gorleku et al., 2011). Our results suggest that palmitoylation of DHHC5 is not required for correct trafficking of DHHC5 to the plasma membrane nor for the interaction between DHHC5 and the endocytic machinery as the endocytic interaction motifs are unaffected by these mutations (Brigidi et al., 2015). The data presented here suggest that when palmitoylated DHHC5 is not bound to Golga7b, it is endocytosed. Preventing palmitoylation of the GluR1 subunit of the AMPA receptor inhibits its endocytosis (Hayashi et al., 2005), which is similar to what we propose for DHHC5. Interestingly, the three palmitoylation sites on the C-terminal tail of DHHC5 map to an amphipathic alpha helix in the corresponding crystal structure of DHHC20 (Rana et al., 2018) and this palmitoylation could potentially pin this helix to the plasma membrane thereby altering the conformation of the C-terminal tail. As the majority of DHHC5 is palmitoylated on two or more sites *in vivo* (Collins et al., 2017), it suggests that palmitoylation could play a major role in the regulation of cell surface expression of DHHC5.

The observation that DHHC5 requires an interaction with Golga7b for stabilisation at the plasma membrane appears to be in conflict with the finding that C-terminal DHHC5 mutant, which does not interact with Golga7b, is stabilised at the plasma membrane. The DHHC5 C-terminal mutant, while being unable to interact with Golga7b, is unable to be internalised because it is not palmitoylated on its C-terminus. This raises the possibility that the interaction with Golga7b functions to protect C-terminally palmitoylated DHHC5 from endocytosis and suggests that palmitoylation of the C-terminal tail of DHHC5 is an endocytic trigger for the protein.

Activity-dependent trafficking of DHHC5 has been described to occur in neurons after long-term potentiation (Brigidi et al., 2015). Here, we have confirmed that DHHC5 is endocytosed from the plasma membrane by the AP2-regulated clathrin mediated endocytosis pathway in HeLa cells. Interestingly, the results presented here suggest that DHHC5 is trafficked as part of its normal lifecycle and does not require stimulation of cells in any way, as DHHC5 is restored to the plasma membrane within 30 minutes of dynasore treatment. This could be a general feature of DHHC5 in all cell types or it could have different cycling dynamics in various specialised cell types. Importantly, inhibition of endocytosis corrects the localisation defect when DHHC5 is co-expressed with mutant Golga7b. This demonstrates that the palmitoylation deficient Golga7b does not prevent insertion of DHHC5 into the plasma membrane. Instead, Golga7b acts to stabilise DHHC5 at the plasma membrane by preventing its internalisation.

Our unbiased AP-MS experiments uncovered a new role for DHHC5 in cell adhesion. Palmitoylation regulates several proteins involved in cell:cell and cell:matrix adhesion (Little et al., 1998; Roberts et al., 2014; Lievens et al., 2016; Roberts et al., 2016; Aramsangtienchai et al., 2017) and for many of them, palmitoylation regulates their localisation. Among cell adhesion complexes, desmosomes are notable for the number of proteins in the complex that are palmitoylated, with at least 6 members of the complex palmitoylated in A431 cells (Roberts et al., 2014), and in this work, we have shown that DHHC5 is the PAT responsible for the palmitoylation of the key complex members desmoglein-2 and plakophilin-3. We have also demonstrated that knockdown of DHHC5 affects the trafficking and localisation of DSG2, which had been demonstrated previously to be dependent on DSG2 palmitoylation, albeit with a noticeably milder phenotype than we observed (Roberts et al., 2016). It is also notable that the knockdown of Golga7b also caused defects in DSG2 localisation and trafficking demonstrating a functional role for Golga7b in cell adhesion. Plasma membrane localisation of DHHC5 is required for the correct trafficking and localisation of DSG2. This implies that plasma membrane localisation of DHHC5 is needed for efficient DSG2 palmitoylation, trafficking and incorporation into desmosomes. However, this appears to contradict the previous work which suggested that palmitoylation of DSG2 occurs at the Golgi which facilitates trafficking to the plasma membrane (Roberts et al., 2016). The reasons for this apparent contradiction are unclear, but the same study showed that palmitoylation-deficient DSG2 is still able to reach the plasma membrane, indicating that palmitoylation of DSG2 is not strictly required for trafficking to the plasma membrane and that once DSG2 reaches the plasma membrane it is palmitoylated by DHHC5 which facilitates its incorporation and therefore stabilisation in desmosomal complexes. However, DHHC5 appears to cycle between the plasma membrane and endomembranes, making is possible that DSG2 is palmitoylated by DHHC5 at non plasma membrane sites, similar to what has been demonstrated for δ-catenin at dendritic spines (Brigidi et al., 2015).

Furthermore, we have determined that plakophilin-3 is also a DHHC5 substrate and as it has been shown previously that palmitoylation of plakophilin-3 regulates the localisation of DSG2 as well as a number of other desmosomal components (Roberts et al., 2014), this would indicate that DHHC5 regulates the localisation and assembly of desmosomes more generally. The finding that plakophilin-3 is a DHHC5 substrate also helps to explain why the localisation and trafficking phenotype of DSG2 upon DHHC5 knockdown is more severe than when DSG2 palmitoylation alone is prevented. Loss of PKP3 palmitoylation has a profound effect on the localisation of DSG2 (Roberts et al., 2016). These findings reveal a completely novel role for DHHC5 in the regulation of desmosome function.

We established a broader role for DHHC5 in cell adhesion by observing increased scattering of DHHC5 depleted cells due to a reduction in cell adhesion. To our knowledge, this is the first report of a specific PAT that regulates the palmitoylation of essential adhesion proteins and that its depletion leads to defects in cell adhesion. The link between DHHC5 and adhesion is not without precedent, however, as a number of known DHHC5 substrates have roles in cell adhesion. Flotillin-2 regulates the stability of desmoglein-3 at the plasma membrane and depletion of Flot2 leads to reduced desmosomal adhesion (Vollner et al., 2016). As DHHC5 is the PAT responsible for Flot2 palmitoylation (Li et al., 2012), it is possible that DHHC5 could have a role in desmosomal adhesion through this pathway in addition to palmitoylation of PKP3 and DSG2 as demonstrated in this work. Another DHHC5 substrate δ-catenin (Brigidi et al., 2014) competes with p120ctn to interact with the adhesion protein E-cadherin. δ-catenin binding to E-cadherin reduces p120ctn at cell junctions and inhibits adhesion and E-cadherin binds to desmogleins 2 and 3 to promote desmosome formation (Shafraz et al., 2018). This suggests that DHHC5 could influence the assembly, stability and location of desmosomes through a number of mechanisms, and this work shows that it has a direct regulatory role for at least two members of the demosomal protein complex.

The regulation of cell adhesion by DHHC5 has some potentially interesting implications for DHHC5 in disease. Disruption of desmosomes in the heart is known to lead to the serious heart condition arrhythmogenic right ventricular cardiomyopathy (ARVC) (Kant et al., 2015). To the best of our knowledge, palmitoylation of desmosomal proteins has not been implicated in ARVC to date, but given that DHHC5 has been shown to be enriched in the heart (Ohno et al., 2006), it is possible that palmitoylation of desmosomal proteins by DHHC5 in the heart could play a role in this condition. It also opens up the possibility of DHHC5 being used as a marker for potential cancer metastasis, as the results presented here suggest that a lower level of DHHC5 in a cell would lead to an increased chance of metastasis due to reduced cell adhesion.

In conclusion, the results presented in this work establish a new mechanism of DHHC5 localisation through the action of Golga7b and implicate DHHC5 and Golga7b as important members of cellular adhesion machinery. Going forward, the exact molecular mechanism of how Golga7b influences DHHC5 localisation and how their interaction is mediated will be important questions to address to fully understand this novel regulatory mechanism. It will also be important to investigate the triggers that control the endocytosis of DHHC5 and how palmitoylation and phosphorylation of DHHC5 work in concert to control its presence at the plasma membrane.

## Materials and methods

### Cell culture

All cells were grown in normal DMEM (Gibco) supplemented with 1% penicillin/sptreptomycin (Gibco), except for during Calcium switch experiments, when A431 cells were grown in calcium free DMEM (Gibco). All transfection was carried out using polyethylinimine (PEI, 25kDa linear, Alfa Aesar) using a ratio of 1:3 of DNA:PEI with cells allowed to grow for 24 hours after transfection before harvesting or fixing unless otherwise stated. For siRNA treatments, siRNA was mixed with mission siRNA transfection siRNA transfection reagent (Sigma Aldrich) and incubated for 30 minutes at room temperature before the mixture was added to cells. SiRNA used: DHHC5, Golga7b, AP2µ, universal non-targeting control (All Dharmacon, SMARTPool)

### Constructs

N-terminally HA-tagged DHHC5 construct was a kind gift from Dr Andrew Peden. All other contructs were ordered from Genscript with either human DHHC5 (GeneID 25921) or human Golga7b (GeneID 401647) as templates. All ORF clones were under the CMV promoter in the pcDNA3.1+ vector which had either a FLAG or HA tag following the protein coding region before the stop codon. SiRNA was obtained from Dharmafect.

### Antibodies

Anti-DHHC5 (Atlas Antibodies), for ICC 1:250, for immunoblotting 1:1000. Anti-Golga7b (Bioss Antibodies), for ICC 1:100, for immunoblotting 1:1000. Anti-FLAG (Sigma Aldrich), for ICC 1:500, for immunoblotting 1:1000. Anti-HA (Sigma Aldrich), for ICC 1:200. Anti-AP2µ (Genetex), for ICC 1:150, for immunoblotting 1:1000. Anti-DSG2 (Santa Cruz Biotech), for ICC 1:200, for immunoblotting 1:250. Anti-PKP3 (Santa Cruz Biotech), for immunoblotting 1:200. Anti-Flotillin-2 (Cell Signalling Technology), for immunoblotting 1:1000. Anti-Actin (Cell Signalling Technology), for immunoblotting 1:2500. Anti-Rabbit HRP (Thermo Fisher Scientific), for immunoblotting 1:2500. Anti-Mouse HRP (Cell Signalling Technology), for immunoblotting 1:2500. Anti-Mouse Alexa Fluor 488 (Invitrogen), for ICC 1:2500. Anti-Rabbit Alexa Fluor 594 (Invitrogen), for ICC 1:2500. Anti-Chicken Alexa Fluor 647 (Invitrogen), for ICC 1:2500.

### Acyl-Biotin Exchange

Cell pellets were collected and resuspended in lysis buffer (2.5% sds, 50mM Tris pH 7.2, 50mM NaCl, 1mM EDTA) and then homogenised using an electric homogeniser for 15 seconds. Lysates were heated at 70°C for 15 minutes before being centrifuged at 16,000xg for 10 minutes and the supernatants retained. After cooling, maleimide (Sigma Aldrich) was added to 100mM final concentration and incubated for 3 hours at 40°C with shaking. Samples were then acetone precipitated by the addition of 4x volume of −20°C acetone and resuspended twice in lysis buffer with 5 washes with ice cold acetone after each precipitation to remove any excess maleimide. Lysates were split into + and – hydroxylamine (HA) treatment conditions. The +HA samples had 2M hydroxylamine/50mM Tris pH 7.4 added to a final hydroxylamine concentration of 1M, while the –HA samples had an equal volume of 50mM Tris pH 7.4 added. HA was made fresh immediately before use and prepared by dissolving hydroxylamine in distilled water before the pH was adjusted up to 7.4 with NaOH. EZ link Biotin-HPDP (Thermo Fisher) dissolved in DMSO, mixed with N,N-dimethylformamide and then added to samples at a final concentration of 0.5mM. Samples were then incubated for 1 hour at 37°C with shaking. Proteins were precipitated with acetone and then washed 4 times with ice cold acetone to remove any excess biotin. Protein was resuspended in IP buffer (1% triton x-100, 50mM tris pH 7.2, 150mM NaCl, 1mM EDTA, protease inhibitor cocktail 1:200 (Sigma)) and incubated with 30µl of streptavidin-sepharose resin (Thermo Fisher), which was prewashed twice with IP buffer, for 1 hour at room temperature with end-over-end rotation. The resin was washed 4 times in the IPbuffer before elution of bound proteins by the addition of 20mM TCEP for 20 minutes at 37°C with shaking. Eluted proteins were collected by centrifugation at 800xg for 5 minutes and the supernatants were retained. Samples were stored at −20°C until further use.

### Immunoprecipitation

Cell pellets were collected and immediately resuspended in lysis buffer (1% triton x-100, 50mM tris pH 7.2, 150mM NaCl, 1mM EDTA, protease inhibitor cocktail 1:200 (Sigma)). Samples were then rotated end-over-end at 4°c for 30 minutes before spinning at 16,000xg for 5 mins. The supernatants were retained. 100µl of anti-FLAG M2 resin (Sigma Aldrich – A2220) was washed in lysis buffer without protease inhibitors 3 times, added to the samples and then left overnight with end-over-end rotation at 4°c. Samples were then spun at 800xg for 2 minutes to collect the resin before being washed 5 times for 5 minutes each at room temperature with end-over-end rotation with lysis buffer without protease inhibitors. Immunoprecipitated proteins were eluted by 3 times resin bed volume of 2mg/ml triple FLAG peptide (Sigma Aldrich) dissolved in lysis buffer without protease inhibitors for 30 mins at room temperature with end over end rotation. The samples were then centrifuged at 800xg for 5 minutes and the supernatants retained. Eluted proteins were stored at −20°C until further use. For immunoprecipitations of endogenous proteins from mouse forebrain, a whole mouse forebrain was added to a lysis buffer (1% triton x-100, 1% sodium deoxycholate, 50mM tris pH 7.2, 150mM NaCl, 1mM EDTA, protease inhibitor cocktail 1:200 (Sigma)) and homogenised 10 times with a dounce homogeniser on ice. Samples were spun at 16,000xg and the supernatant retained as above. Samples were pre-cleared with protein G sepharose before centrifugation and retention of the supernatant. Samples were then incubated with either anti-DHHC5 or anti-Golga7b antibodies and protein G sepharose overnight at 4°C. The resin was then washed 5 times with lysis buffer before elution of bound proteins by heating the resin for 20 minutes at 70°C.

### Immunocytochemistry

Cells were seeded onto glass coverslips in a 12 well plate and were grown for 24 hours. The media was changed and cells were transfected with a 3:1 ratio of PEI:cDNA for 24 hours as described previously. Media was removed, coverslips were washed briefly with 1-2 ml of PBS and cells were fixed with 4% PFA for 10 minutes at room temperature. Cover slips were washed again with PBS before permeabilisation with 0.3% triton x-100-PBS (vol/vol) for 15 minutes at room temperature with gentle agitation. After washing with PBS, coverslips were blocked with 0.2% fish skin gelatin (Sigma Aldrich)/0.01% triton x-100 (vol/vol) in PBS (blocking solution) for an hour at room temperature with agitation. Primary antibody was incubated at the concentrations mentioned previously diluted in blocking solution for 1 hour at room temperature. Coverslips were washed three times for 5 minutes with 1ml of PBS with gentle agitation before incubation with secondary antibodies diluted in blocking solution for 2 hours at room temperature, protected from light by wrapping plates in tin foil with gentle agitation. Coverslips were then washed with 1ml of PBS 3 times for 5 minutes with gentle agitation before mounting on glass slides with DAPI fluoromount-G mounting media (Southern Biotech) before being kept in dark at 4°C before confocal imaging.

### Calcium switch assays

A431 cells were seeded on glass coverslips in a 12 well plate with either normal DMEM or low Calcium DMEM and placed back into the incubator for 18 hours. Control or targeted siRNA was then added as described previously and cells were placed back into the incubator for 72 hours. CaCl_2_ was then added to the low calcium media to a final concentration of 2mM for 30 minutes, 1 or 3 hours. Control cells were left in low calcium media. Cells were then fixed with 4% PFA and prepared as described previously in the immunocytochemistry section.

### Confocal Microscopy

All confocal microscopy was performed on a Nikon A1 confocal microscope using the following laser wavelengths: 405nm (blue), 488nm (green), 561nm (red) and 640nm (far-red). Images were acquired using a 60x objective lens and the NIS elements program (Nikon instruments) in an ND2 format. For all samples, no primary and no secondary controls were used to threshold the exposure at each individual laser wavelength. All images were analysed using the FIJI (Schindelin et al., 2012) and imageJ (Schneider et al., 2012) software packages. To assess plasma membrane or internal protein signal, the colour channels were split to isolate the channel of interest. The free draw tool was used to draw around a single cell and the mean intensity, raw integrated density and integrated density were measured. The shape was then moved just within the plasma membrane of the cell and the same measurements recorded. The ratio of the two means was then used to calculate the level of protein signal at the plasma membrane (higher ratio indicates more signal at the plasma membrane).

### Immunoblotting

Samples were run on 10% tris-glycine acrylamide gels unless otherwise stated and proteins were transferred to 0.2µm polyvinylidene difluoride membranes (Thermo Fisher) for 90 minutes at 30V Membranes were blocked for 1 hour at room temperature on a roller in pbs 0.1%-tween-20 (vol/vol, pbs-t) 5% skim milk powder (weight/vol), before overnight incubation with primary antibody diluted in pbs-t 1% skim milk powder (weight/vol) at 4°C on a roller. Membranes were then washed 5 times with 10 ml pbs-t for 5 minutes each on a roller at room temperature, before incubation with secondary antibody diluted in pbs-t 1% milk for 1 hour at room temperature. Membranes were washed again 5 times with 10ml pbs-t for 5 minutes before a final wash with 10ml pbs. Clarity ECL solution (Bio-Rad) was added in a 1:1 ratio of peroxide solution:luminol/enhancer solution with a final volume of 10ml for 5 minutes at room temperature on a roller before another wash with pbs on a roller at room temperature. Blots were imaged by exposure on CL-X posure x-ray film (Thermo Fisher) prior to development to visualise protein bands.

### Mass spectrometry

In-gel digestion was performed as reported previously (Bayes et al., 2017). Extracted peptides were re-suspended in 0.5% formic acid and analysed by nanoflow LC-MS/MS using an Orbitrap Elite (Thermo Fisher) hybrid mass spectrometer equipped with a nanospray source, coupled to an Ultimate RSLCnano LC System (Dionex). The system was controlled by Xcalibur 2.1 (Thermo Fisher) and DCMSLink 2.08 (Dionex). Peptides were desalted on-line using a micro-Precolumn cartridge (C18 Pepmap 100, LC Packings) and then separated using a 130 minute gradient from 3%-40% buffer B (0.5% formic acid in 80% acetonitrile) on an EASY-Spray column, 50 cm × 50 µm ID, PepMap C18, 2 µm particles, 100 Å pore size (Thermo). The LTQ-Orbitrap Elite was operated with a cycle of one MS (in the Orbitrap) acquired at a resolution of 60,000 at m/z 400, with the top 20 most abundant multiply-charged (2 + and higher) ions in a given chromatographic window subjected to MS/MS fragmentation in the linear ion trap. An FTMS target value of 1e6 and an ion trap MSn target value of 1e4 was used and with the lock mass (445.120025) enabled. Maximum FTMS scan accumulation time of 500 ms and maximum ion trap MSn scan accumulation time of 100 ms were used. Dynamic exclusion was enabled with a repeat duration of 45 s with an exclusion list of 500 and an exclusion duration of 30 s. The mass spectrometry proteomics data have been deposited to the ProteomeXchange Consortium via the PRIDE [1] partner repository with the dataset identifier PXD011553

### Mass spectrometry data analysis

All raw mass spectrometry data was analysed with MaxQuant version 1.5.6 (Cox & Mann, 2008). Data were searched against a human UniProt sequence databases (downloaded June 2015) using following search parameters: digestion set to Trypsin/P, methionine oxidation and N-terminal protein acetylation as variable modifications, cysteine carbamidomethylation as a fixed modification, match between runs was enabled with a match time window of 0.7 minutes and a 20 minute alignment time window, label free quantification was enabled with a minimum ratio count of 2, minimum number of neighbours of 3 and an average number of neighbours of 6. PSM and protein match thresholds were set at 0.1ppm. A protein FDR of 0.01 and a peptide FDR of 0.01 was used for identification level cut offs. For immunoprecipitation data, protein groups file generated by MaxQuant were loaded into the Perseus platform (Tyanova et al., 2016) with the LFQ intensity from each individual run as main columns. The matrix was filtered to remove all proteins that were potential contaminants, only identified by site and reverse sequences. The LFQ intensities were then transformed by log2(x) and individual LFQ columns were grouped by experiment (e.g. all control experiments grouped, all DHHC5 pulldowns grouped etc.). Rows without valid values in 3 out of 4 runs in at least one of the groups were removed and all remaining missing values were replaced from the normal distribution with each column imputed individually. Following this, a two-tailed Student’s T-test with S0 set to 3 and a permutation based FDR of 0.05 was performed and all non-significant proteins were removed and the matrix was exported in a .txt format for further analysis. For further filtering of results, files analysed using the online Contaminant Repository for Affinity Purification Mass Spectrometry Data program (Mellacheruvu et al., 2013) with additional controls (CC130, CC135, CC136, CC137). Proteins with an SP score above 0.95 were considered bona fide interactors. Gene ontology analysis was performed using the WebGestalt platform (Zhang et al., 2005) with the following parameters: species set to homo sapiens, reference gene set was “genome protein coding”, FDR set to 0.01 with Benjani Hochberg multiple test adjustment.

### Cell Scatter Assay

10,000 A431 cells were seeded in single wells of 6 well plates 24 hours after treatment with either DHHC5 or non-targeting siRNA. Cells were grown for 24 hours before media was removed and replaced with serum free DMEM and cells were incubated for 8 hours. Cells were imaged at 10x magnification on a Nikon Widefield microscope system, positions of fields of view recorded, EGF was added to a final concentration of 5ng/ml and cells were incubated for 48 hours. Cells were then imaged again at the same positions as previously. To quantify scattering, the grid tool of ImageJ was used, with the number of grid squares occupied by cells compared at 0 and 48 hours.

### Trans-epithelial electrical resistance

125,000 A431 cells were seeded in transwell inserts in 24 well plates which had a 0.4µm pore size. Cells were allowed to grow for 12 hours until confluency before treatment with siRNA, which was performed for 24 hours. The resistance in each well was measured 3 times with an epithelial voltohmmeter. The measurements for each well were averaged and a blank value of a transwell insert without cells was subtracted from each measurement before statistical analysis.

## Acknowledgements

M.O.C is supported by grants from the BBSRC (BB/P021689/1 & BB/R003491/1). K. T. W was funded by a University of Sheffield scholarship. We would like to thank Darren Robinson from Wolfson Light Microscopy Facility and Adelina Acosta Martin from the biOMICS mass spectrometry facility for providing technical assistance. We would also like to thank Andrew Wood and Dr Kai Erdmann for their assistance with the TEER experiments and Dr Andrew Peden for useful discussions and critical reading of the manuscript.

## Author contributions

K.T.W and M.O.C conceived the experiments, K.T.W performed experiments and data analysis and K.T.W and M.O.C wrote the paper.

