## Supplemental Information for "S-acylated Golga7b stabilises DHHC5 at the plasma membrane to regulate desmosome assembly and cell adhesion"

**Supplemental Figure 1: Efficiency of protein depletion with siRNAs.** (A) Immunoblot from HeLa cells treated with 10nM DHHC5 siRNA (right lane) or non-targeting siRNA (left lane). Quantification normalised to actin signal. (B) Immunoblot from HeLa cells treated with 20nM Golga7b siRNA (right lane) or non-targeting siRNA (left lane). Quantification normalised to actin signal. (C) Immunoblot from HeLa cells treated with 15nM AP2m siRNA (right lane) or non-targeting siRNA (left lane). Quantification normalised to actin signal.

**Supplemental Figure 2: Overexpression of DHHC5 causes a localisation defect that is rescued by prevention of palmitoylation of the C-terminus of DHHC5.** (A) Confocal images of endogenous DHHC5 in HeLa cells. (B) Confocal images of HeLa cells over-expressing WT DHHC5 C-terminally tagged with a FLAG epitope. (C) Confocal images of HeLa cells over-expressing a C-terminally FLAG tagged catalytic mutant of DHHC5 where the cysteine of the catalytic tetrad of DHHC5 has been replaced with a serine. (D) Confocal microscopy images of HeLa cells over-expressing DHHC5 with the three palmitoylated cysteines in the C-terminal tail mutated to serine residues. (E) Quantification of membrane signal of DHHC5 seen in A-D.

**Supplemental Figure 3: Calcium switch of A431 cells treated with non-targeting siRNA**. (A) Time zero after calcium chloride addition. (B) 30 minutes after calcium chloride addition. (C) 1 hour after calcium chloride addition. (D) 3 hours after calcium chloride addition. (E) Percentage of DSG2 signal seen at the plasma membrane at each time point, *=$p<0.05$, unpaired t test.

**Supplemental Figure 4: Calcium switch of A431 cells treated with DHHC5 siRNA.** (A) Time zero after calcium chloride addition. (B) 1 hour after calcium chloride addition. (C) 3 hours after calcium chloride addition. (D) percentage of DSG2 signal seen at the plasma membrane at each time point

**Supplemental Table S1: DHHC5/WT Golga7b AP-MS data.** Proteins found as significantly enriched in DHHC5 pulldowns when it was co-express with WT Golga7b from HeLa cell lysates, filtered with our own controls plus 4 controls from CRAPome. All proteins with a SAINT probability (SP) of above 0.95 were considered interactors.

**Supplemental Table S2: DHHC5/mutant Golga7b AP-MS data.** Proteins found as significantly enriched in DHHC5 pulldowns when it was co-express with mutant Golga7b from HeLa cell lysates, filtered with our own controls plus 4 controls from CRAPome. All proteins with a SAINT probability (SP) of above 0.95 were considered interactors.

**Supplemental Table S3: DHHC5 AP-MS data.** Proteins found as significantly enriched in DHHC5 pulldowns from HeLa cell lysates, filtered with our own controls plus 4 controls from CRAPome. All proteins with a SAINT probability (SP) of above 0.95 were considered interactors.

### Supplementary Figure 1

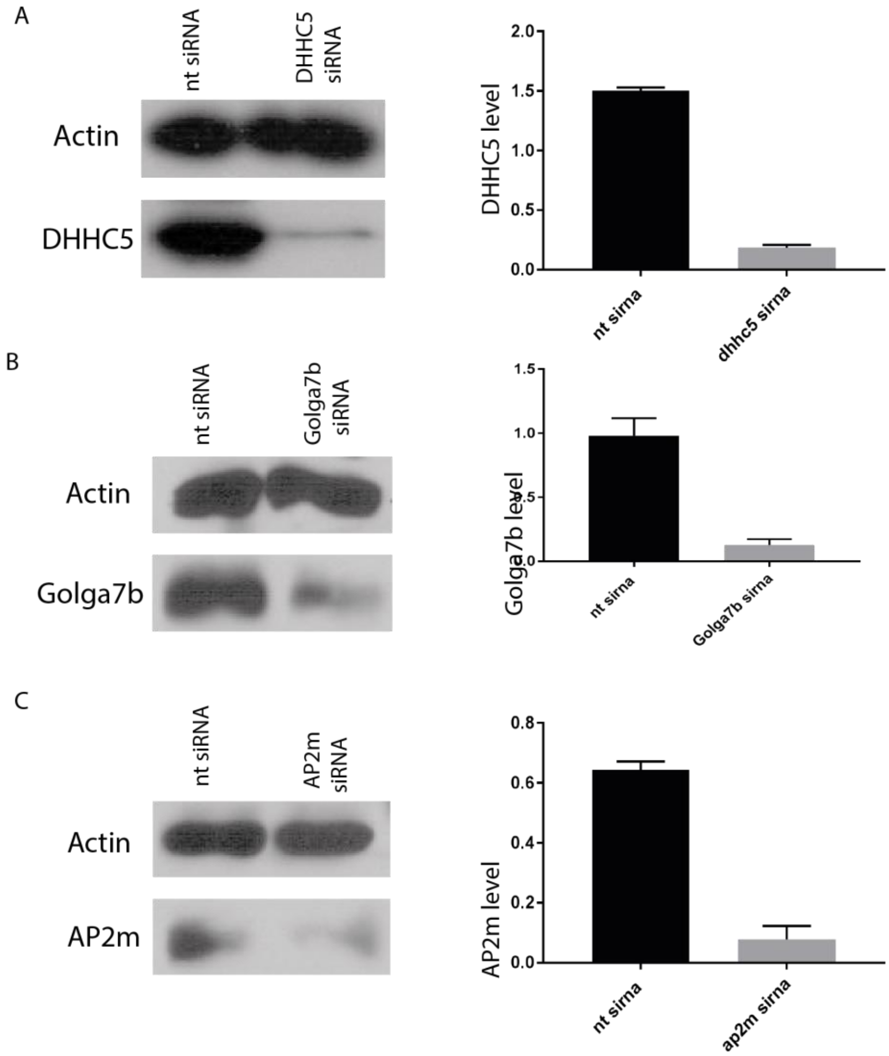

### Supplementary Figure 2

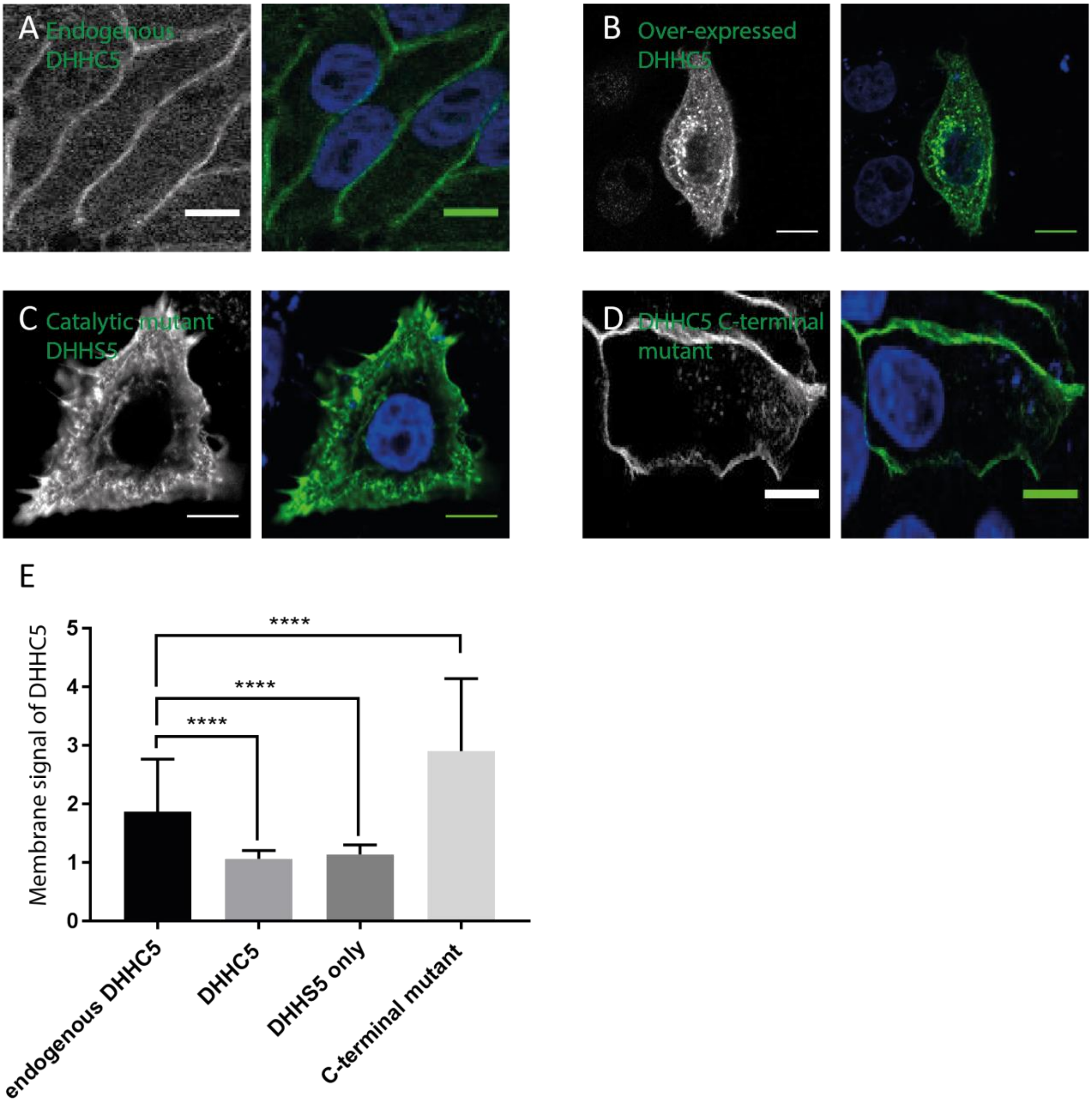

### Supplementary Figure 3

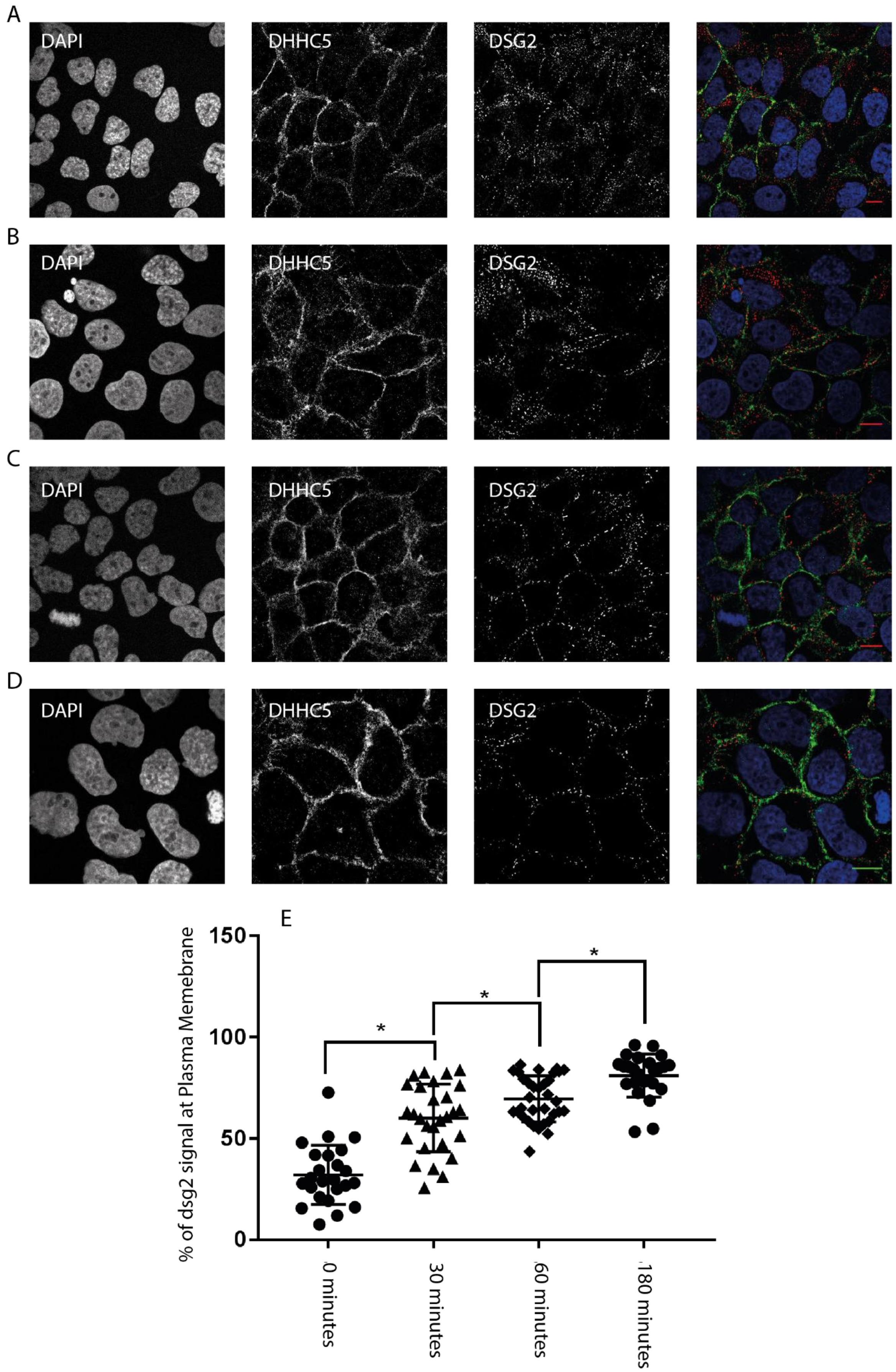

### Supplementary Figure 4

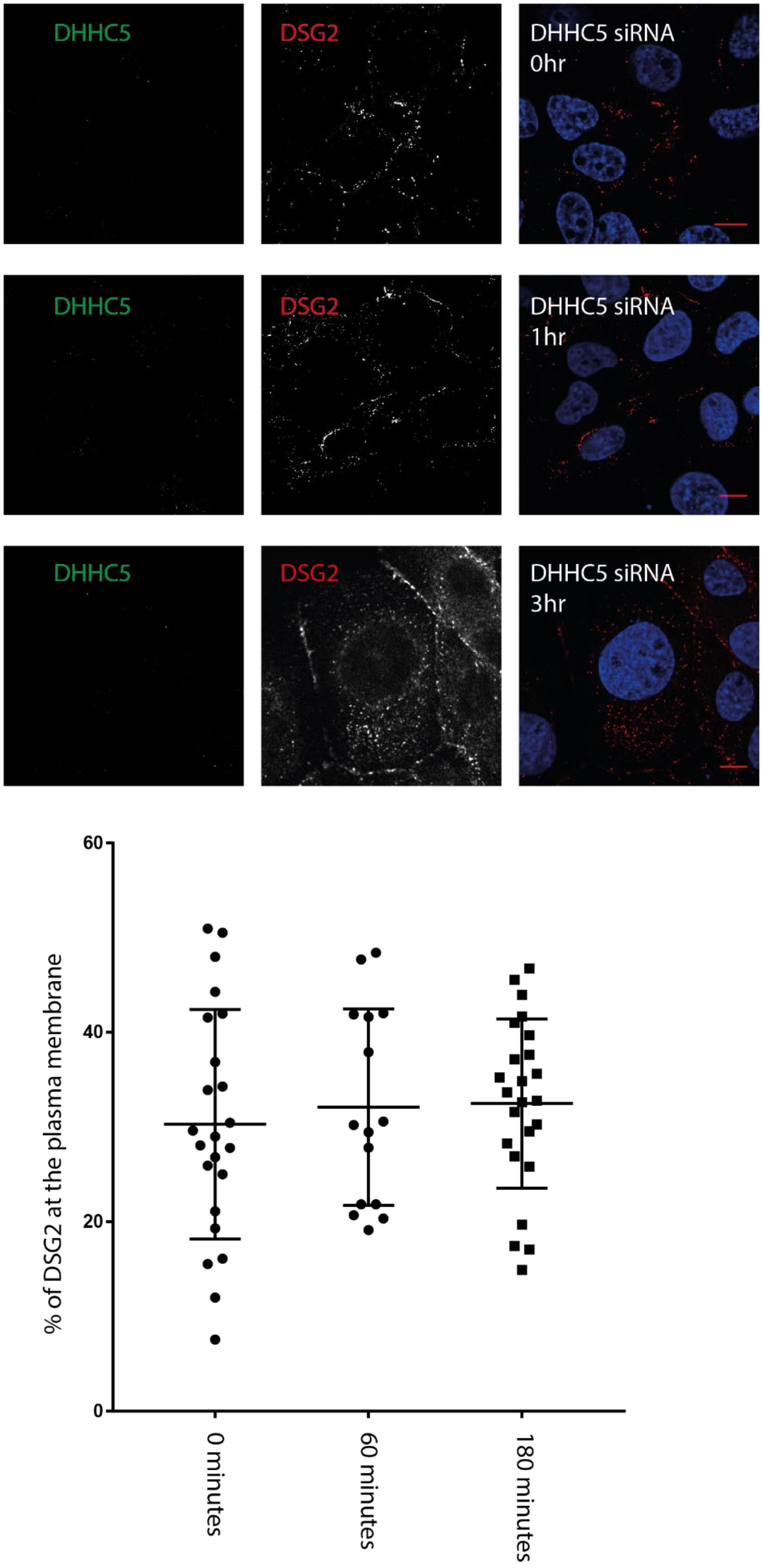
